# Computational enhancement of single-cell sequences for inferring tumor evolution

**DOI:** 10.1101/341743

**Authors:** Sayaka Miura, Louise A Huuki, Tiffany Buturla, Tracy Vu, Karen Gomez, Sudhir Kumar

## Abstract

**Motivation**: Tumor sequencing has entered an exciting phase with the advent of single-cell techniques that are revolutionizing the assessment of single nucleotide variation (SNV) at the highest cellular resolution. However, state-of-the-art single-cell sequencing technologies produce data with many missing bases (MBs) and incorrect base designations that lead to false-positive (FP) and false-negative (FN) detection of somatic mutations. While computational methods are available to make biological inferences in the presence of these errors, the accuracy of the imputed MBs and corrected FPs and FNs remains unknown.

**Results**: Using computer simulated datasets, we assessed the robustness performance of four existing methods (OncoNEM, SCG, SCITE, and SiFit) and one new method (BEAM). BEAM is a Bayesian evolution-aware method that improves the quality of single-cell sequences by using the intrinsic evolutionary information in the single-cell data in a molecular phylogenetic framework. Overall, BEAM and SCITE performed the best. Most of the methods imputed MBs with high accuracy, but effective detection and correction of FPs and FNs require sampling a large number of SNVs. Analysis of an empirical dataset shows that computational methods can improve both the quality of tumor single-cell sequences and their utility for biological inference.

**Conclusions**: Tumor cells descend from pre-existing cells, which creates evolutionary continuity in single-cell sequencing datasets. This information enables BEAM and other methods to correctly impute missing data and incorrect base assignments, but correction of FPs and FNs remains challenging when the number of SNVs sampled is small relative to the number of cells sequenced.

**Availability:** BEAM is available on the web at https://github.com/SayakaMiura/BEAM.

## 1 Introduction

Tumor sequencing is yielding critical insights into somatic drivers of tumorigenesis and clonal structure of heterogeneous tumors (Brastianos, et al., 2015; Gawad, et al., 2014; Gundem, et al., 2015; McFadden, et al., 2014; Nassar, et al., 2015; Navin, et al., 2011; Nik-Zainal, et al., 2012; Sanborn, et al., 2015; Xue, et al., 2017; Yachida, et al., 2010; Zhao, et al., 2016). The rapid advancement of single-cell sequencing technologies has made it possible to profile somatic mutations carried by individual cells (Eirew, et al., 2015; Francis, et al., 2014; Gawad, et al., 2014; Gawad, et al., 2016; Huang, et al., 2015; Hughes, et al., 2014; Navin, 2014; Navin, 2015; Paguirigan, et al., 2015; Shapiro, et al., 2013; Van Loo and Voet, 2014; Yu, et al., 2014; Zafar, et al., 2016). Many studies have performed single-cell sequencing on tumors to identify clones and their evolutionary relationships (Eirew, et al., 2015; Gawad, et al., 2014; Hou, et al., 2012; Jan, et al., 2012; Li, et al., 2012; Melchor, et al., 2014; Navin, 2015; Potter, et al., 2013; Xu, et al., 2012; Yu, et al., 2014). Thus, single-cell sequencing will be instrumental in revealing the genetic changes that occur during cancer progression, which is a prerequisite for clone identification and the inference of evolutionary relationships among cells and relative timing of mutation events. But, the utility of current single-cell sequencing technologies is limited by many technical issues (Gawad, et al., 2016; Navin, 2014; Navin, 2015; Ning, et al., 2014; Wang and Navin, 2015). For example, the low physical coverage of some genomic regions and positions prevents unambiguous assignment of a nucleotide base to those positions, known as “missing bases” (MBs). Allelic dropout (ADO) events cause false-negatives (FNs) when mutant alleles are present but not amplified. Infidelity of amplification can cause false-positives (FPs) when errors during initial amplification are inherited to subsequent molecules and a “mutation” is identified that was not present in the sampled cell. Sometimes a single-cell cannot be completely separated from other cells, which results in the sequencing of multiple cells together. FP rates (3×10^−5^ – 7×10^−5^ per homozygous wild-type positions) and ADO rates (0.2 – 0.4 per heterozygous site) can exceed the rate of occurrence of true mutations (Ross and Markowetz, 2016). MBs also occur at frequencies as high as 58% (Hou, et al., 2012) in single-cell data sequences (Gawad, et al., 2016). All of these problems result in inaccurate single-cell sequences even when high sequencing coverage has been achieved.

Many new methods have been developed to compensate for these issues and allow reliable inference from single-cell sequence datasets. For example, OncoNEM (Ross and Markowetz, 2016) and BitPhylogeny (Yuan, et al., 2015) identify clones and their evolutionary relationship (i.e., clone phylogeny). SiFit infers cell phylogeny with the consideration of sequencing errors (Zafar, et al., 2017). SCITE (Jahn, et al., 2016) and Kim and Simon (2014) methods are designed to infer the order of mutation accumulated over time in a tumor from the single-cell sequences. SCG is designed to deal with the issue of multi-cell sequencing when inferring clone sequences (Roth, et al., 2016).

These methods produce corrected single-cell sequences, but they do not report their performance in imputing MBs correctly and reducing FPs and FNs. The primary focus of these current methods has been to improve the quality of biological inferences from error-containing single-cell sequencing data. Consequently, the absolute and relative performance of current methods for reducing the error present in single-cell sequences is not known. Significant improvement in the quality of single-cell sequences will enable use of a large number of sophisticated methods in molecular phylogenetics (Nei and Kumar, 2000) for inferring the evolutionary history of clones, reconstructing ancestral clones, identifying early and late occurring driver mutations, and characterizing inter- and intra-tumor heterogeneity. These standard approaches cannot currently be used for tumor single-cell data, because they are not robust to the presence of high levels of sequence error (Zafar, et al. (2017). For example, a widely-used maximum likelihood method (Stamatakis, 2014) produces a cell phylogeny (**Fig. 1b**) from simulated single-cell sequence data (with MBs, FPs, and FNs) that is clearly very different from the true tree (**Fig. 1a**). In addition to various inconsistencies in the evolutionary relationships, the branch lengths leading to the tips of the phylogeny are extensively overestimated, because all the cells of a clone (same color) are actually identical (**Fig. 1a**). Consequently, the inferred cell phylogeny shows much greater evolutionary depth, resulting in inflated estimates of tumor heterogeneity and incorrect mapping of mutations. Therefore, single-cell sequences require correction before use in downstream biological analysis.

**Figure 1.**
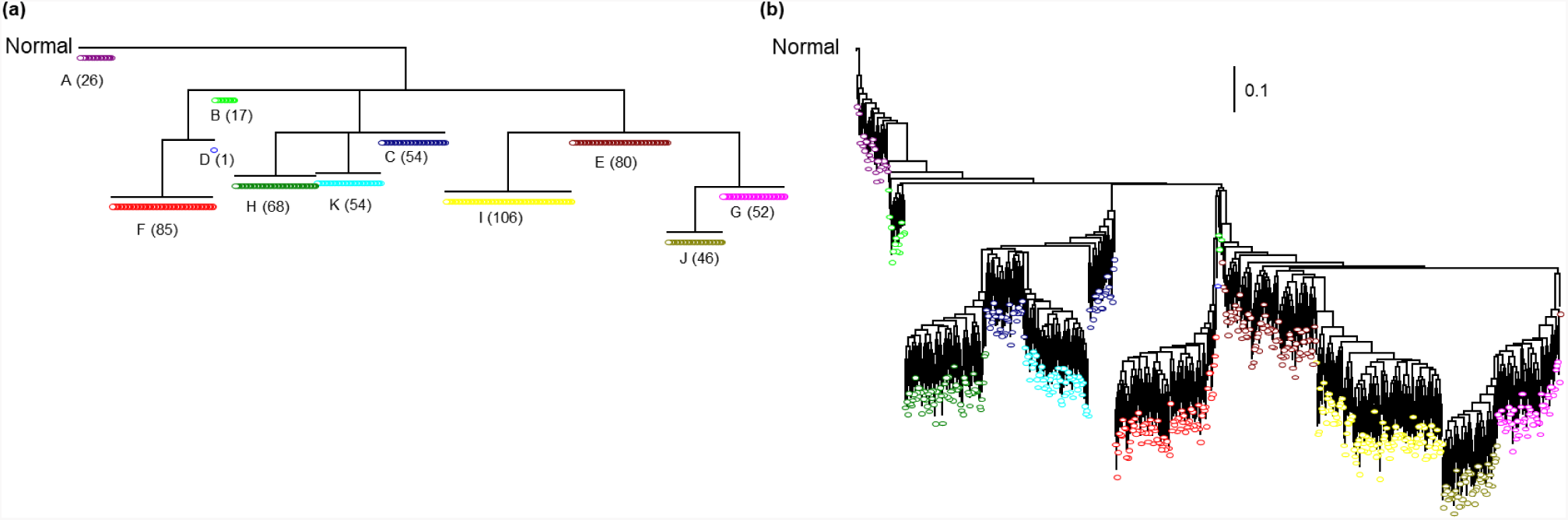
Impact of missing data and sequencing errors on the inferred cell phylogeny. (**a**) The true evolutionary tree of 1,000 cells distributed among tumor clones A-K (shown in different colors); the number of cells sampled for each clone is in parentheses. (**b**) Cell phylogeny inferred using simulated single-cell sequences in which 500 SNVs were sampled. Roth, et al. (2016) software and parameter settings were used to generate the data with 20% missing bases, 28% false-positives, and 7% false-negatives. A maximum likelihood method for phylogenetic analysis of SNV data (Stamatakis, 2014) was used to infer the cell phylogeny. Branch lengths are drawn to scale (number of SNVs/site). The inferred cell phylogeny shows greater sequence divergence than the true phylogeny due to the influence of many false-positive mutations.

In this article, we present the performance of four existing methods (OncoNEM, SCG, SCITE, and SiFit) in correctly imputing MBs and reducing the numbers of FPs and FNs. We excluded methods that did not produce single-cell sequences, e.g., BitPhylogeny (Yuan, et al., 2015) and the method of Kim and Simon (2014). In addition, we propose and test a new method, Bayesian Evolution-Aware Method (BEAM), which employs molecular phylogenetics and a Bayesian prediction framework to improve the quality of single-cell sequences (see **Methods**). Our testing focused on computer simulated datasets, as knowledge of the true single-cell sequences enables direct assessment of the performance of computational methods (Ross and Markowetz, 2016; Roth, et al., 2016). We also analyzed one empirical dataset (Li, et al., 2012) to gauge the utility of computational approaches in a real-world scenario and the concordance of the inferences produced.

In the following, we present information on the simulated data used in our evaluation of methods, followed by a description of the BEAM approach and the assumptions, parameters, and accuracy measures used. We then present results from our analyses of simulated and empirical data discuss the patterns observed.

## 2 Methods

### 2.1 Generation of datasets by computer simulations

<u>Roth et al. datasets (R1000×50 and R100×50 datasets):</u> We used the simulator and parameter settings described by Roth, et al. (2016) to produce 240 datasets. This simulator first generates a clone phylogeny and then the clone genotypes. A new model phylogeny is generated for every dataset (e.g., **Fig. 1a** for 1,000 cells and 10 distinct clones). To generate a clone phylogeny, new clones are created by accumulating mutations (a mutation rate set to 0.1 per site) until all SNV loci (50) are created. The simulator uses an infinite sites assumption, so no mutations override each other, and introduces loss of heterozygosity at a rate of 0.2 per site in which heterozygous mutants are changed into homozygous mutants without allowing any loss of mutations. A clone genotype is assigned to each cell by sampling from a categorical distribution (clonal prevalence), which is generated from a symmetric Dirichlet distribution with the parameter value of 1. Datasets with 100 and 1,000 cells were produced by Roth, et al. (2016) in which doublet single-cell sequencing was simulated by sampling two clone genotypes at different rates: 5%, 10%, 20%, and 40% of the cells. The simulator generated allelic count data with ADO by sampling from the empirical distribution of SNV frequencies specified in Roth, et al. (2016). The depth of coverage at each locus was chosen from a Poisson distribution with a mean of 1,000 reads. The number of variant reads was sampled from a Binomial distribution with the parameter selected from an empirical distribution and the depth of coverage sampled from a Poisson distribution. To determine if an allele was present or absent, the Binomial exact test was performed and a p-value threshold of 10^−6^ was used. In the resulting data, 3–51% of the observed mutant alleles were FPs and 2–20% of the observed homozygous wild-type alleles were FNs. We randomly assigned a “missing” value (MB) to 20% of the bases.

<u>Ross and Markowetz datasets</u> (M10×50 – M50×300 datasets). This collection of 690 datasets was generated using the Ross and Markowetz (2016) simulator and parameter settings. In their approach, clone phylogenies were first generated by iteratively adding a branch with a node to an existing node that was randomly chosen from a growing phylogeny (1, 5, 10, and 20 clones). Unobserved clones were then introduced by removing clones that had at least two descendant clones (0, 1, 2, 3, and 4 unobserved clones). A new clone phylogeny was generated for each dataset, and each cell is assigned to a clone with a probability corresponding to its size (10, 20, 30, and 50 cells). **Figure 2a** shows an example phylogeny of 20 cells used to simulate clone evolution for generating M datasets (M10×50 – M50×300 datasets) Along the clone phylogeny, true clone genotypes were generated by assigning mutations with a uniform probability (50, 100, 200, and 300 SNVs). Observed genotypes were derived from true genotypes by introducing MBs (10%, 20%, 30%, and 40% of SNVs), FPs (10^−5^%, 5%, 10%, 20%, and 30% of mutant alleles), and FNs (5%, 10%, 20%, and 30% of wild-type alleles).

**Figure 2:**
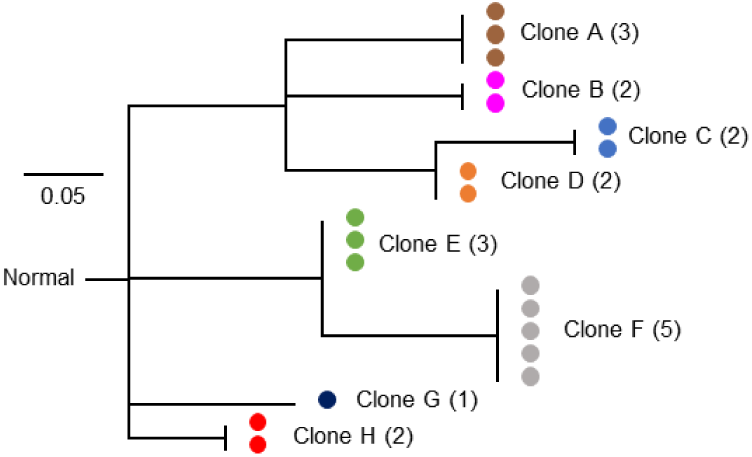
An example cell phylogeny used in computer simulations. In this phylogeny, there are eight distinct clones (A − H), each represented by 1 – 5 cells. This phylogeny was used to generate one dataset by Ross and Markowetz (201 6) simulator.

### 2.2 Accuracy measurements

We recorded the numbers of missing bases (MBs), false-positives (FPs), and false-negatives (FNs) in the simulated single-cell sequence datasets. The total number of correct positions (with no MB, FP, or FN) were aggregated and divided by the product of the number of cells and the number of SNVs. This quantity is referred to as the initial sequence quality (*Q*_0_), which is the same as the mean Hamming distance between the true and the inferred single-cell sequence. After the sequence data was subjected to computational analysis by BEAM, OncoNEM, SCG, SCITE, and SiFit, the sequence quality was reassessed (*Q*_BEAM_, *Q*_OncoNEM_, *Q*_SCG_, *Q*_SCITE_, and *Q*_SiFi_, respectively).

While the positions containing MBs, FPs, and FNs are known for simulated data, no such information exists in the analysis of empirical data and the computational methods must be applied to all the positions. Therefore, we also compared the total numbers of MBs, FPs, and FNs before and after the application of a computational method to a dataset.

### 2.3 New method evaluated (BEAM)

The new Bayesian evolution-aware method (BEAM) uses classical molecular evolutionary phylogenetics to impute missing data and detect base assignment errors in the single-cell sequencing data. It is based on the premise that significant evolutionary information is present in the initial cell sequences regardless of base assignment errors. For example, cells from the same clone show a strong tendency to occur in close proximity in the initial cell phylogeny (**Fig. 1b**), as seen by the location of cells marked by the same color in the true tree (**Fig. 1a**), despite the presence of a large number of MBs, FPs, and FNs in the simulated sequence data. BEAM uses this intrinsic evolutionary information and computes a Bayesian posterior probability (PP) of observing all possible alleles at each SNV position in each single-cell sequence, as described below.

For brevity, we explain BEAM using an example dataset that was generated using the cell phylogeny in **Figure 2a**. It consists of 20 single-cells from eight distinct clones and 200 SNVs. The simulated sequence dataset contained 800 MBs, 429 FPs, and 106 FNs (**Fig. 3a**). For this data, we first infer a cell phylogeny from the observed single-cell sequences by using a maximum likelihood method specifically suited for phylogenetic analysis of SNV data (Stamatakis, 2014) (**Fig. 3a**). This approach does not require the infinite sites assumption, i.e., mutations are allowed to be lost and they may occur at the same genomic position in different cells, which is different from the principle applied in OncoNEM, SiFit, and SCITE (Jahn, et al., 2016; Ross and Markowetz, 2016; Zafar, et al., 2017).

**Figure 3.**
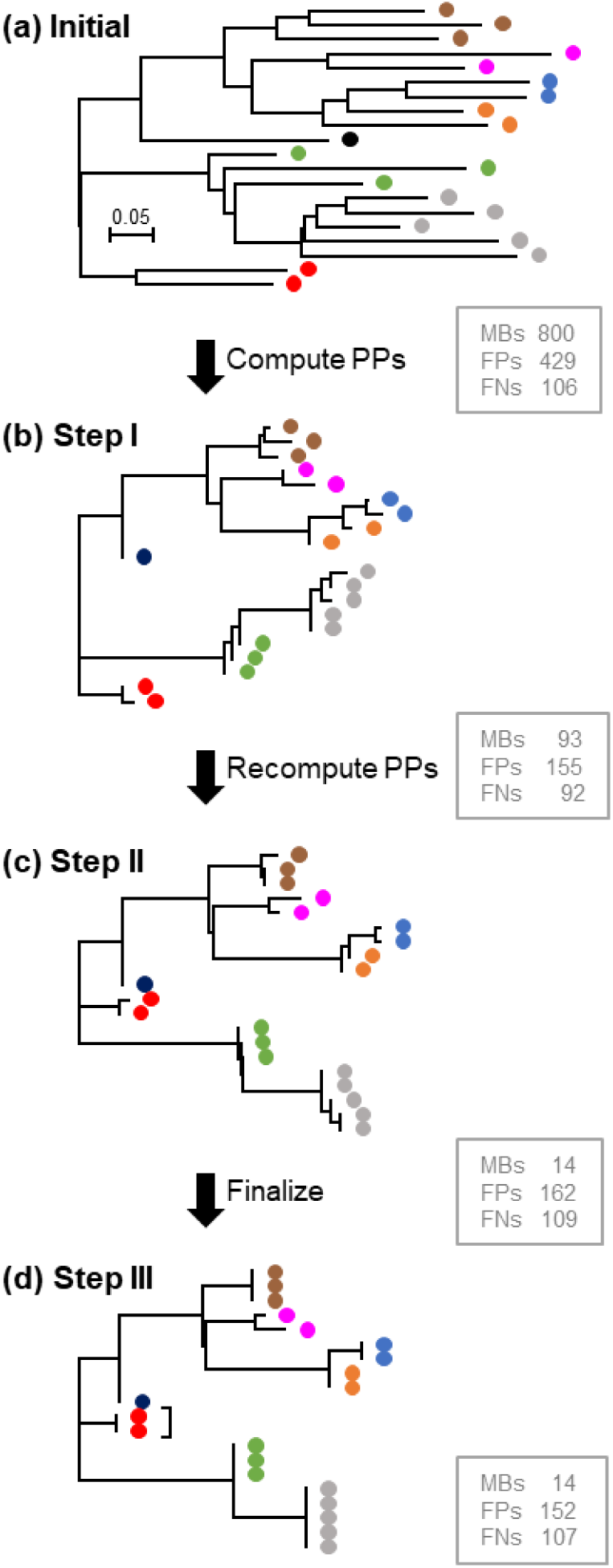
An overview of the BEAM approach. (**a**) The initial cell phylogeny, along with branch lengths, derived from single-cell sequence data simulated using the model tree in **Figure 2**. The Ross and Markowetz (2016) simulator and parameter settings were used. The number of MBs, FPs, and FNs in the initial sequence data are shown in the box. (**b**) Cell phylogeny produced from the data in which MBs were imputed and FPs and FNs were corrected. The remaining MBs, FPs, and FNs are shown. (**c**) Improved cell phylogeny after recomputing PPs by using the phylogeny in panel **b**. (**d**) The final cell phylogeny produced by BEAM along with the remaining MBs, FPs and FNs. The topology and branch lengths are very similar to those in the model tree shown in **Figure 2**.

In this example, cells from the same clones (the same color) generally cluster together, but identical cells of a clone can show extensive observed sequence divergence (e.g., brown cells in **Fig. 2a** and **3a**). Given this initial cell phylogeny and the initial cell sequences, we estimate PP of each possible base assignment at each position in a cell sequence. This computation is explained by considering a set of four sequences (**Fig. 4**). In this tree, *x*_1_ to *x*_3_ represent the nucleotides at a given position in the tumor cell sequence; *x*_4_ is the wild-type base from the normal cell sequence.

**Figure 4.**
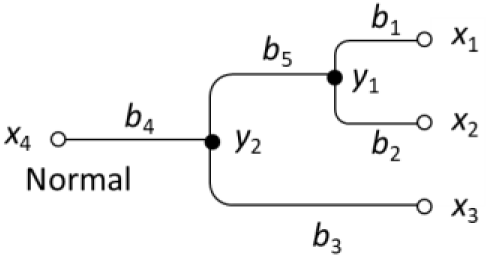
A simple tree used in explaining PP computation. *x_i_* is a base in the single-cell sequence, *y_i_* is a base in the ancestral sequence, and *b_i_* refers to a branch length.

We estimate relative probabilities of different bases to assign to *x*_1_. We represent nucleotides at the three other tip nodes by the vector ***x*** = (*x*_2_, *x*_3_, *x*_4_) and let ***y*** = (*y*_1_, *y*_2_) represent the vector of nucleotides *y*_1_ and *y*_2_ at the two ancestral nodes in this phylogeny. In order to estimate *x*_1_ given the single-cell sequences 1 to 4, we compute the Bayesian posterior probability for each possible set of nucleotides (*x*_1_, ***y***) following Liu, et al. (2016):
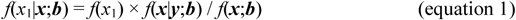

Here, ***b*** is the vector of branch lengths *b*_1_, …, *b*_5_. We compute the probability *f*(*x*_1_|***x*;*b***) for all possible combinations of *x*_1_, *y*_1_, and *y*_2_. The posterior probability (PP) of base A at *x*_1_, for example, is given by:
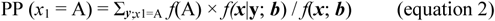

We compute PP for each possible base assignment at each position (*i*) in the single-cell sequence 1. If there are *m* single-cell sequences and *n* SNV positions in the dataset, *m*×*n* computations are performed. We assigned the base with PP > 0.7 to the position of interest. This cut-off was selected because it led to the best performance in computer simulations. When none of the bases show PP > 0.7 at a position, we assign the base or missing value originally observed at that position.

In our example, BEAM improves the single cell sequences by imputing base identities at MB positions by reducing these from 800 to 93, reducing FPs from 429 to 155, and reducing FNs from 106 to 92. Because of these improvements, the inferred cell phylogeny becomes more accurate (**Fig. 3b**). At the same time, the erroneously long tip branches in the initial phylogeny are shortened via elimination of FPs (**Fig. 3a**).

In the next step, we use the new cell phylogeny (**Fig. 3b**) and recompute PPs by applying equations 1 and 2. Now, MBs decrease from 93 to 14 and FPs and FNs increased slightly (155 to 162 and 92 to 109, respectively; **Fig. 3c**). The cell phylogeny inferred using these single-cell sequences looks similar to the true phylogeny (**Fig. 2**).

The next step is clone annotation. Clones delineation can proceed by using the bootstrap procedure to assess the robustness of the branching patterns (Felsenstein, 1985; Nei and Kumar, 2000), merging cells by iterative clustering of nodes along branches (Ross and Markowetz, 2016), or by using a k-medoids clustering approach on a distance matrix that is obtained from the latest cell phylogeny (Zafar, et al., 2017). BEAM provides the user with an option to assign identical sequences to all the cells of a clone by erasing potentially spurious mutations. In our simulations, we found that it was best to assign identical clone sequences to cells that were connected with effectively zero branch lengths in the phylogeny when the number of new mutations was small (default: <2% of SNVs; **Fig. 3d**). In this case, BEAM will finalize sequences by computing PPs of each possible base assignment and comparing the average PPs between potential bases in order to assign the base with the highest average PP to all the cells from the same clone. Ultimately, BEAM produces refined single-cell sequences as well as the cell phylogeny.

### 2.4 Options used for analyzing data

We used default or recommended parameters to perform SCG, SCITE, OncoNEM, and BEAM analyses. SCG (Roth, et al., 2016) was performed with the doublet option. When the status of mutations (presence or absence) within a predicted clone genotype had <0.95 probability, we assigned missing values to those positions. To assign a clone for each cell, SCG additionally computed the probability of having a predicted clone genotype for each cell. Thus, we assigned the predicted clone genotype with the highest probability. When none of predicted clone genotypes had a high probability (>0.01) for a cell, no predicted clone genotype was assigned to the cell. Cells lacking clone genotype assignments were removed, and datasets consisting entirely of cells lacking clone genotype assignments were removed from accuracy considerations as we considered that SCG failed to correct sequences for these datasets.

OncoNEM (Ross and Markowetz, 2016) analyses were performed using true rates of false positive and false negative base assignment errors in the input files (observed sequences). Maximum Bayes factor for which a smaller model was preferred was 10, and the model search stopped when the best scoring tree stabilized for at least 200 iterations. Mutant nucleotides were assigned when the probability of observing the mutation at a given position was > 0.95 and missing bases were assigned when the probability of observing the mutant nucleotide was between 0.05 and 0.95. SCITE (Jahn, et al., 2016) analyses were performed by giving true rates of false-positive and false-negative detections of mutations. The desired number of repetitions of the MCMC was 1, and the desired chain length of each MCMC repetition was 900,000. Often, mutant nucleotides were not assigned to any cells. When mutant nucleotides were not assigned to any cells, those sites were assigned with wild-type bases. When multiple possible cell phylogenies for a dataset were produced, we used the option to marginalize out the alternatives. We distinguished heterozygous and homozygous mutations for R1000×50 and R100×50 datasets. When multiple cell sequences were inferred for a single cell, we replaced inconsistent base assignments with MBs. For SiFit (Zafar, et al., 2017) analyses, we also input true rates of false positive and false negative detection of mutations. The number of iterations run for each restart was 10,000. Cell genotypes were inferred by inferring the order of mutations along the cell phylogeny predicted by SiFit. LOH rate and deletion rate were set to zero.

### 2.4 Empirical data

We analyzed a single-cell sequencing dataset from muscle-invasive bladder tumors (Li, et al., 2012), which has been analyzed previously in other articles proposing new methods (Ross and Markowetz, 2016; Zafar, et al., 2017). We obtained sequenced reads in FASTQ format by using SRA toolkit (v2.8.1)(Leinonen, et al., 2011). We mapped these sequenced reads to the human genome sequence (hg18 from UCSC database; https://genome.ucsc.edu/) by using the Burrows-Wheeler alignment tool (BWA v0.7.12)(Li and Durbin, 2009) with *aln* and *samse* options. Samtools (v1.3.1)(Li, et al., 2009) was used to remove reads with low mapping quality (≤40) when creating BAM files, which were sorted by chromosome coordinate. This initial data processing follows the protocol described in Zafar, et al. (2016).

We then used Monovar (Zafar, et al., 2016) to call mutations. We performed *mpileup* in Samtools with the options presented in the instructions for Monovar (i.e., minimum base quality was zero). Monovar analysis was performed with the default or recommended options, i.e., offset for prior probability of false-positive error was 0.002; offset for prior probability of allelic drop out was 0.2; threshold for variant calling was 0.05; and the number of threads used in multiprocessing was 2.

For downstream computational analysis, we selected SNVs in coding regions that were identified by Zafar, et al. (2016). Nucleotides identified among the majority of normal cells were assigned as wild-types, and the other bases found in tumor cells were assigned as mutants. Our analyses did not distinguish homozygous and heterozygous mutations. Following Zafar, et al. (2016), we assigned missing values to positions with coverage depth less than 6x, in addition to positions where Monovar did not predict a genotype. Lastly, we removed one cell that contained a very large number of missing bases.

## 3 Results

### 3.1 Analysis of simulated large datasets

We first present results from the analysis of 120 large datasets that consisted of 1000 cells each with 50 SNVs (R1000×50). The initial sequence quality (*Q*_0_) of these datasets ranged from 65−75% (**Fig. 5a**), which was caused by 20% MBs, 7−51% FPs, and 2−16% FNs. Four of the five methods were able to handle these large datasets and produced refined single-cell sequences of much higher quality than the input (**Fig. 5a**). The average *Q*_BEAM_ was 89% and varied from 80% - 95%. The performance was the best when the input sequences contained the fewest errors and the worst when the input sequence error was the largest. SCITE performed similarly well (*Q*_SCITE_ = 90%) and showed very similar trends describing the relationship between input and output quality. While SCG also worked well (*Q*_SCG_ = 89%), it showed much greater variability in performance. SiFit showed the smallest improvements (*Q*_SiFit_ = 80%). Overall, the sequence quality improved for all datasets after a computational method was used.

**Figure 5:**
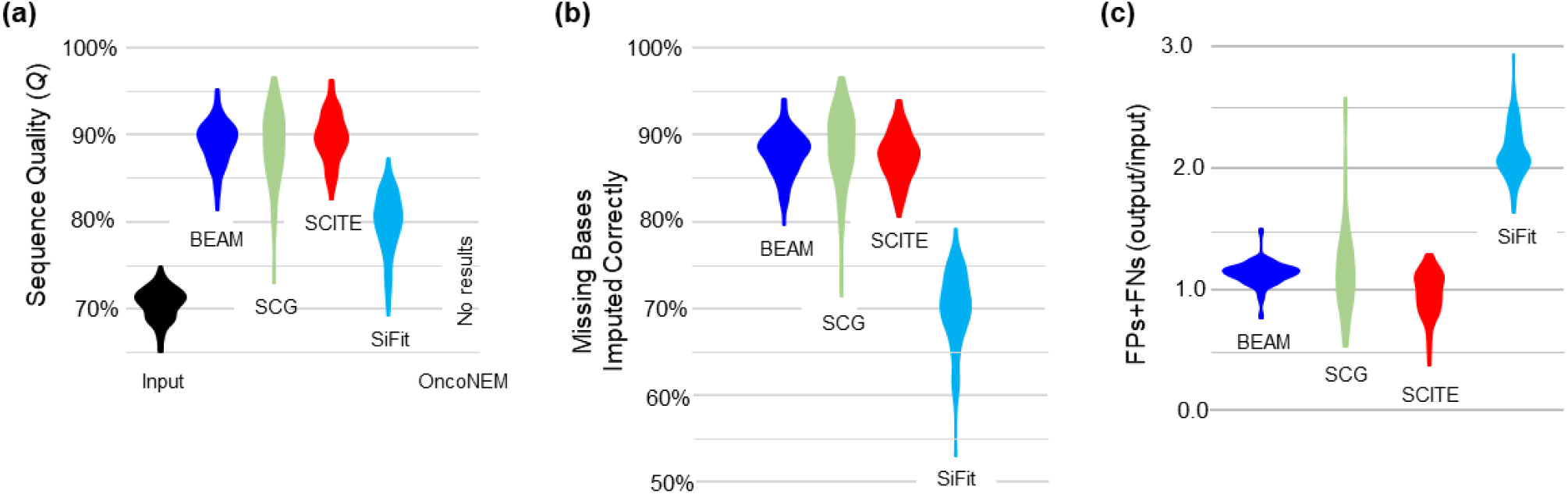
Improvement in the single-cell sequences realized by using computational methods for processing larger datasets (R1000×50). (a) Sequence quality of data input to computational methods and of data output from four computational methods. (b) Proportions of missing bases imputed correctly by computational methods. (c) The ratio of the number of FPs and FNs in the output and the input sequences (output/input).

We found that the correct imputation of MBs was the primary reason for the improvements observed for R1000×50 datasets. The fraction of MBs found in the single-cell sequences decreased, on average, by 100%, 100%, 100%, and 96% after the application of BEAM, SCG, and SCITE, respectively, and most of the missing data were correctly imputed (88%, 88%, and 87%, respectively)(**Fig. 5b**). Although SiFit imputed all of the MBs without producing new MBs, SiFit correctly imputed a much smaller fraction of MBs than the other methods (70% of MBs were correctly imputed). Interestingly, however, no method showed a significant ability to reduce the total number FPs and FNs in this data (**Fig. 5c**), as the numbers of FPs and FNs in the output were greater than in the input (output/input ratio > 1.0). Of all the methods, SiFit showed the worst average ratio (2.18), which can happen because the correction procedure has to be applied to all the bases in the input sequence data as the positions with FP and FN are not known in the real world data analysis. This causes the creation of many new false positives and false negatives.

Next, we tested the accuracy of computational methods for datasets in which the number of sequenced cells was reduced to 100 (R100×50). Again, all methods successfully improved single-cell sequences with very similar output sequence quality (**Fig. 6a**). Outcomes were similar to those observed for R1000×50 datasets, except that SiFit performed much better and OncoNEM produced results. As with the larger dataset (R1000×50), the correct imputation of MBs was the primary improvement observed (**Fig. 6b**), and all of the methods produced sequences in which the numbers of FPs and FNs were similar to or much larger than the error in the input sequence data (**Fig. 6c**).

**Figure 6:**
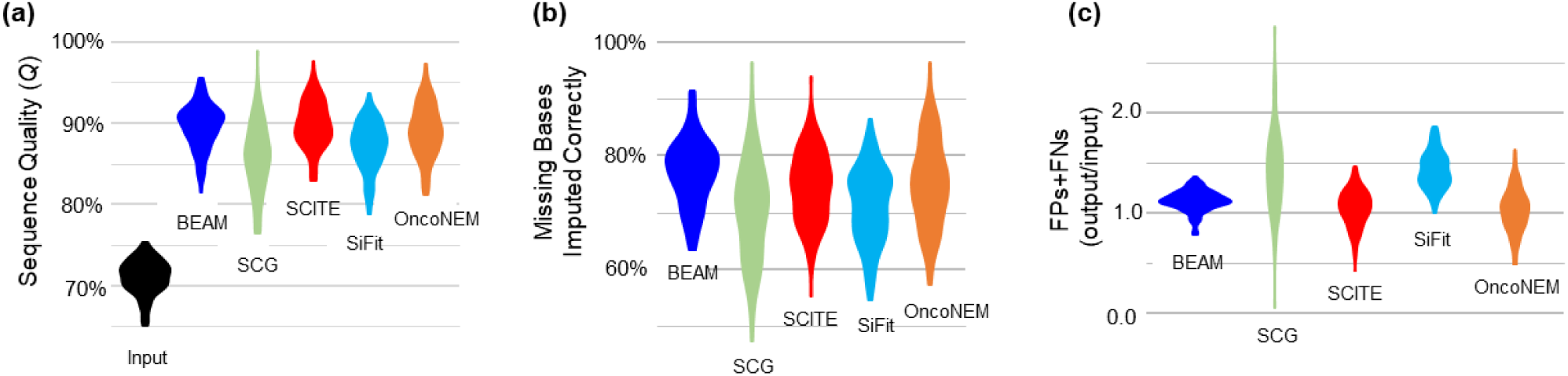
Improvement in the single-cell sequences for medium sized datasets (R100×50). (**a**) Sequence quality of the data input to five computational methods and of output from computational processing. (**b**) Proportions of missing bases imputed correctly by computational methods. (**c**) The ratio of the number of FPs and FNs in output and input sequences (output/input).

In both R1000×50 and R100×50 datasets, we observed that the identification of FPs and/or FNs was less effective than the imputation of MBs for all the methods (**Fig. 5c** and **6c**). We hypothesized that, unlike the imputation of MBs, detection and correction of FPs and FNs was very sensitive to the available cell relationship information that can be gleaned from the initial error-prone single-cell sequencing data. We tested this hypothesis by increasing the number of SNVs used to 500 (R1000×500), while keeping the number of errors per SNV the same as in R1000×50 dataset. We applied BEAM and SCG to the new collection of datasets and found that the output/input ratio of FPs and FNs for BEAM became much less than 1 (**Fig. 7a**). That is, BEAM was able to produce sequences with ~40% fewer FPs and FNs. This improvement over R1000×50 datasets is explained by the fact that the initial cell relationships derived using the input data in BEAM are more accurate when the number of SNVs analyzed is large (compare phylogenies in **Fig. 7c** and **1b**). This improvement enables the Bayesian analysis to generate better predictions, because the phylogenetic prior is closer to the truth.

**Figure 7:**
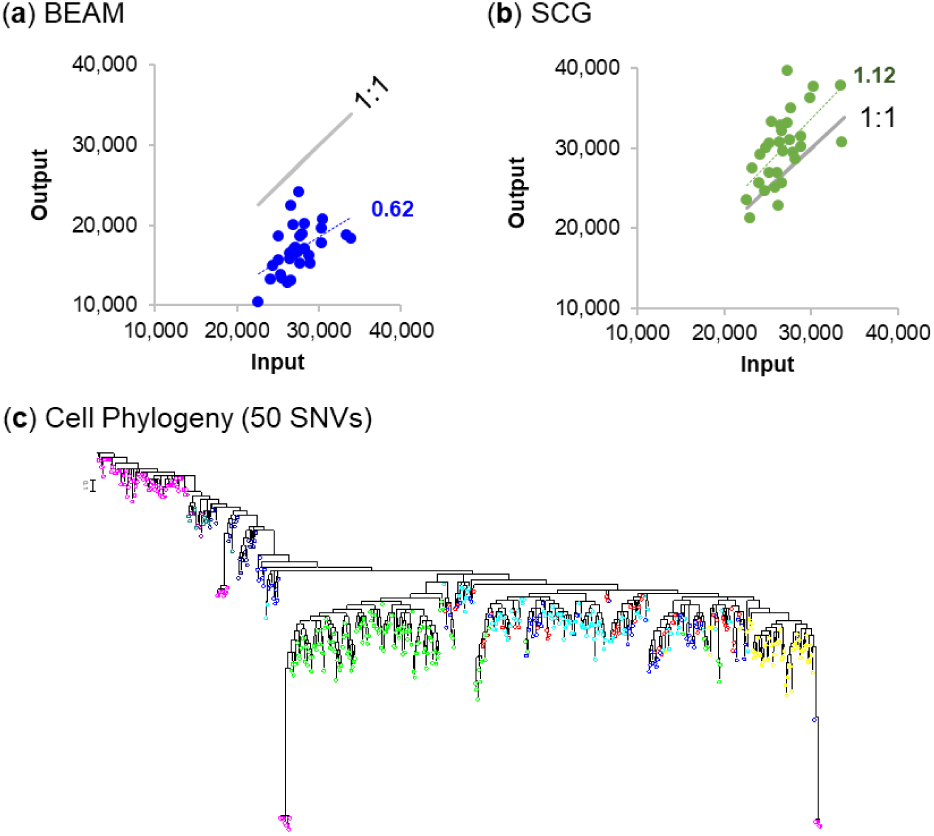
Detection and correction of FP and FNs in datasets containing 1000 cell sequences with SNVs. Scatter graphs show the relationships of the number of the FPs and FNs in the input data (x-axis) and the output data (y-axis) produced by (**a**) BEAM and (**b**) SCG. Linear regression slopes through the origin are 0.62 and 1. 12, respectively. (**c**) The initial cell phylogeny when the number of SNVs is small (50), which is very different from the true cell phylogeny and the cell phylogeny derived using 500 SNVs (**Fig. 1b**).

The performance of SCG did not improve, and still created more FP and FN errors in the output (**Fig. 7b**). This may be because SCG first clusters cells to identify clones and then assigns clone genotypes. This procedure does not appear to benefit from increased evolutionary information in the dataset. Additional analyses support this reasoning, as BEAM, OncoNEM, SCITE, and SiFit were able to detect and correct many FPs and FNs when the number of SNVs sampled was large in relation to the number of cells sampled.

### 3.2 Analysis of smaller datasets

Next, we next analyzed 690 datasets that were generated by Ross and Markowetz (2016) (M datasets). All contained fewer cells than the R1000 datasets (largest) and R100 datasets (medium-sized), but the M datasets contained the same or larger numbers of SNVs (50, 100, 200, and 300 SNVs) compared to the R datasets. All five methods produced results for all datasets, in which the initial quality of cell sequences was between 65 and 75%. BEAM, SCITE, and SiFit increased the quality of the sequences to an average 92% (**Fig. 8a**), but OncoNEM (82%) was less accurate and SCG performed poorly (65%). OncoNEM and SCG did not impute missing data as accurately as BEAM, SCITE and SiFit (**Fig. 8b**).

**Figure 8:**
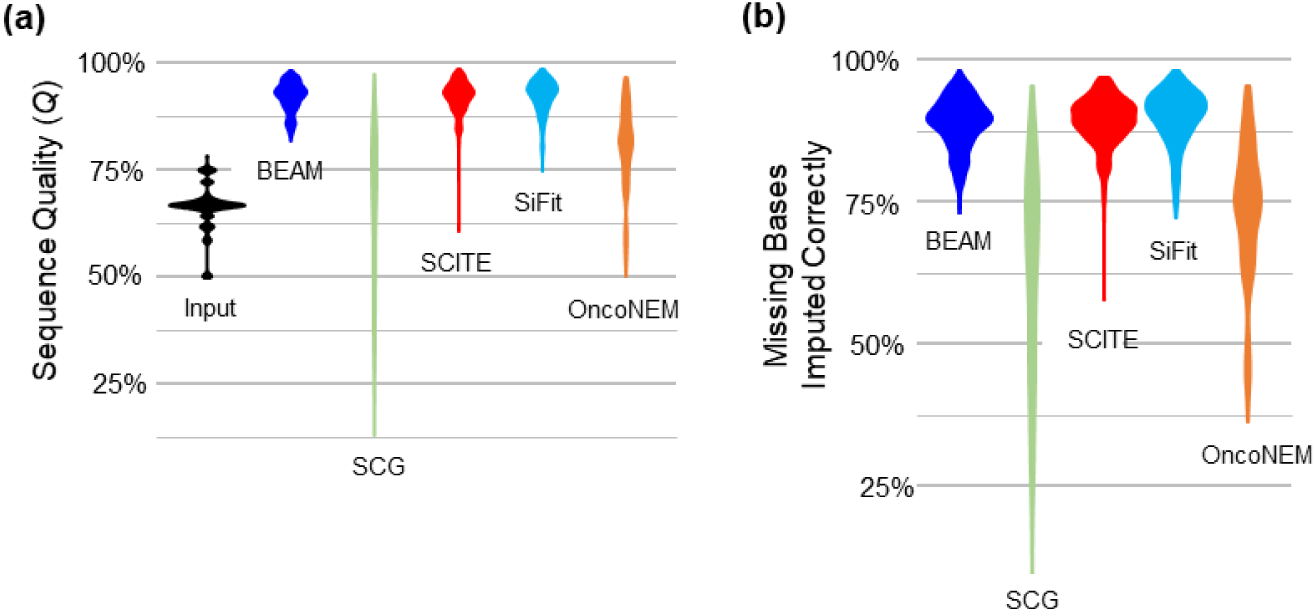
Improvement in the single-cell sequences for datasets containing a small number of cells (M). (**a**) Sequence quality of the data input to computational methods and of the data output from all five methods. (**b**) Proportions of missing bases imputed correctly. These are aggregate patterns from the analysis of 690 datasets; see **figures 9** and **10** for more detailed results.

All methods, except SCG, decreased the numbers of FPs and FNs in the analysis of M datasets (**Fig. 9**), because the ratio of the number of SNVs to the number of cells was greater than that for R datasets. In fact, increasing the number of SNVs provides a proportional increase in the performance of BEAM, OncoNEM, and SiFit. Patterns observed for M datasets (**Fig. 9)** confirm the results for R datasets with 500 SNVs (**Fig. 7)**: BEAM becomes more accurate with larger numbers of SNVs and SCG’s performance does not improve. OncoNEM performed the best in correcting FPs and FNs in M datasets, but achieves this at the expense of producing many MBs (**Fig. 8**).

**Figure 9:**
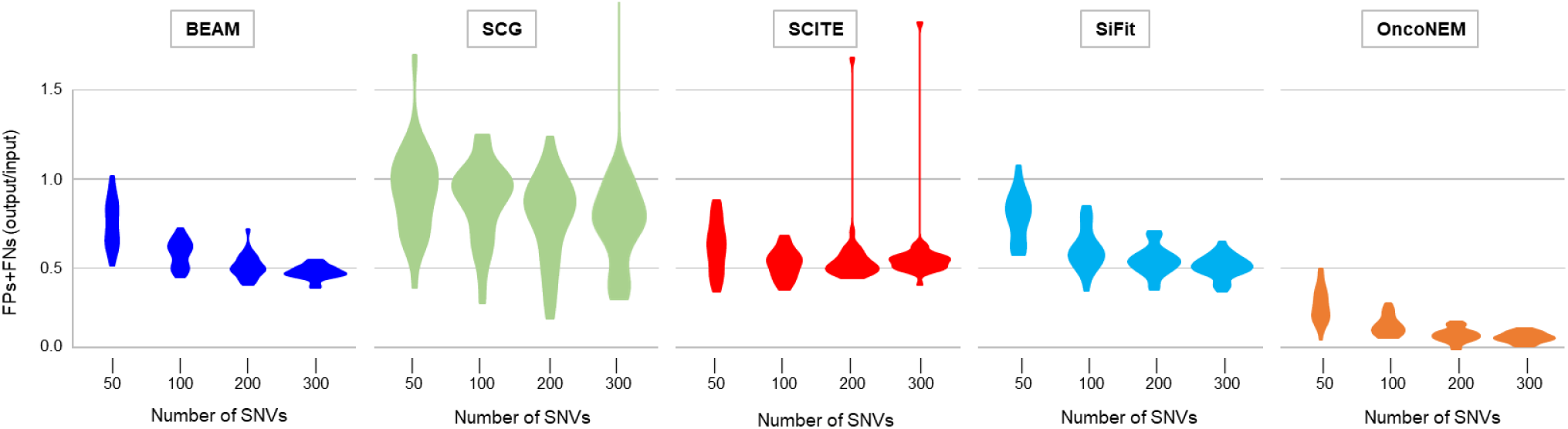
Improving FPs and FNs in the single-cell sequences. The ratio of FP and FN base assignments in the output and the input sequences (output/input) for BEAM, SCG, SCITE, SiFit, and OncoNEM. For SCG, the trend was truncated at 2.0 for simplicity.

As expected, the quality of the inferred sequences became higher as the number of SNVs increased (**Fig. 10a**). The quality of the inferred sequences was higher when the number of cells was high (**Fig. 10b**), because all methods utilize similar cell sequences (or clusters of SNVs) to impute MBs and correct FPs and FNs. For example, the PP calculation is affected by the base assignments of neighboring cells, which will become more accurate when a base assignment is supported by larger number of cells from the same clone. Overall, BEAM, SCITE, and SiFit provided the most robust results when the number of cells was small, and OncoNEM and SCG were greatly impacted if multiple cells per clone were not sampled. Lastly, the quality of the output sequences was a direct function of the fraction of MBs (**Fig. 10c**), FPs (**Fig. 10d**), and FNs (**Fig. 10e**).

**Figure 10:**
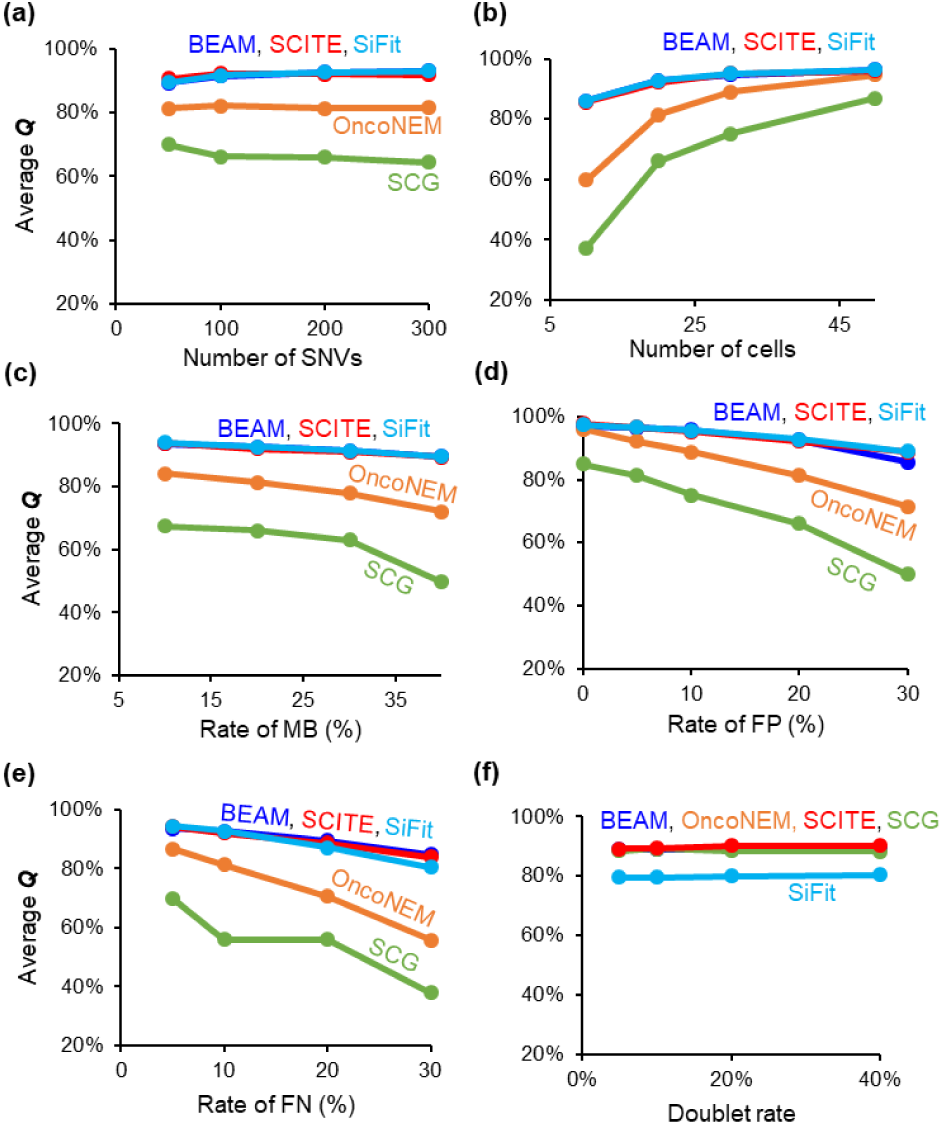
The relationship of the output sequence quality with the various simulation parameters. (**a**) Number of SNVs in the data, (**b**) number of cells sequenced, and fractions of (**c**) missing bases, MBs, (**d**) false positives, FPs, and (**e**) false negatives, FNs in M datasets. (**f**) Effect of increasing doublet sequencing rates on the quality of the output sequencing for R datasets.

Before proceeding with an analysis of empirical datasets, it is important to note that we did not introduce any loss of mutant alleles in our simulations, e.g., loss of heterozygosity (LOH) and the loss of genomic segments. Such mutations will negatively impact the performance of most methods, as the evolutionary relatedness of sequences will be disturbed by such losses. Therefore, such positions should be detected and removed before applying these computational methods. Also, we found that a high rate of doublet sequencing did not adversely impact the performance of any of the computational methods (**Fig. 10f**). This result is consistent with the finding in Jahn, et al. (2016), who reported that doublet sequencing did not lead to lower accuracy in biological inference, which was a major motivation for the development of SCG (Roth, et al., 2016).

### 3.3 Analysis of an empirical dataset

We applied all five methods to a previously published dataset of a muscle-invasive bladder tumor (Li, et al., 2012). This dataset contained 55 cells with 84 SNVs in protein coding regions (see **Methods**). The cell phylogeny before and after the application of BEAM is shown in **Figures 11a** and **11b**, respectively. As observed from computer simulated datasets, the phylogeny based on the initial single-cell sequences shows high diversity among cells, with no clear demarcation of clones; note that we colored tips in **Figure 11a** based on BEAM’s clone predictions in **Figure 11b**). Following the application of BEAM, the cell phylogeny shows distinct clonal structure with 11 different tumor clones (**Fig. 11b**).

**Figure 11:**
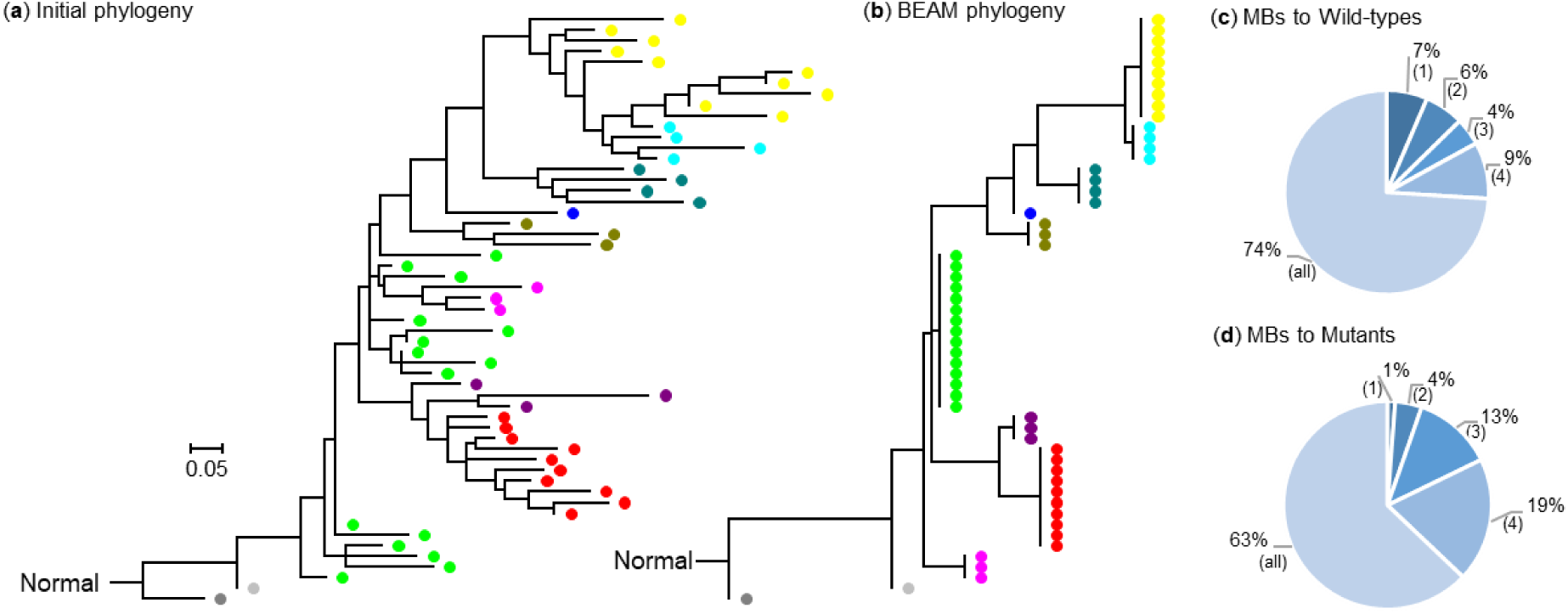
Analysis of an empirical dataset. (**a**) The cell phylogeny produced by using the initial single-cell sequences from Li et al. (2012). (**b**) The cell phylogeny produced by BEAM after imputing MBs and correcting FPs and FNs. The proportions of imputed MBs that were assigned a (**c**) wild-type base and (**d**) a mutant base are shown, with different slices indicating the number of methods (in parentheses) that made the same assignment, including BEAM.

Li et al. (2012) suggested that all of the tumor cells were derived from a single ancestral cell. This conclusion is supported by BEAM, as the inferred cell phylogeny showed that all of the tumor cells had a common ancestral cell. Li et al. (2012) reported that a set of 15 cells, which they identified as a single clone, arose early in the tumor’s evolution. BEAM found that 14 of these cells belonged to a group of early-emerging clones (gray, green, and pink clones in **Fig. 11b**), and one cell was a part of a later arising (yellow) clone. Two BEAM-identified clones (purple and red) are the closest relatives and were contained in the second clone in Li et al. (2012). Cells from the third clone in Li et al. (2012) are divided among 5 closely-related clones by BEAM. Therefore, the cell phylogeny produced by BEAM provides a more detailed clonal structure, which remains consistent with the clonal structure (three major clones) presented by Li et al. (2012).

The initial cell phylogeny (**Fig. 11a**) was transformed into the final cell phylogeny (**Fig. 11b**) by BEAM, because the final single-cell sequences contained only 6 positions with missing values. This is a major improvement, as the original dataset contained 666 positions with missing values. In imputing MBs, BEAM assigned a mutant base to 337 positions and the normal base to 323 positions. 63% of the mutant base and 74% of the wild-type base assignments made by BEAM were also suggested by all other methods (**Fig. 11c** and **d**, respectively). In fact, more than 93% of BEAM’s assignments for MBs were shared with at least one other computational tool. Therefore, extensive consensus exists among methods. We also examined whether BEAM’s base assignments for MBs were supported by the read count data. As mentioned in an earlier section, Monovar does not assign a base when there are only a few reads. So, we tested whether the wild type base assignments by BEAM were supported by higher read counts than other bases at the examined positions. This was indeed the case, the read counts of wild type alleles assignments averaged 7 times higher. Assignment of mutant alleles to MBs were also supported by 22% larger read counts than the wild type alleles. Therefore, read count data generally supports the assignments made by BEAM.

In addition to imputing almost all MBs in the observed sequences (99% of MBs), BEAM corrected wrong mutant and wild-type base assignments. In these predictions, 105 mutation calls were found to be false positives, 65% of which were also detected by at least two other methods. That is, they were consensus assignments (≥ 3 out of 5 methods). Also, 183 wild-types were detected to be false negatives, i.e., they should have been mutant assignments. 88% of these were consensus assignments, becase at least two other methods suggested the same base assignment. Therefore, we expect that the use of these computational methods will enable better biological conclusions.

## 4 Discussion

We have reported that computational methods are generally capable of imputing missing bases with high accuracy and, thus, can improve the quality of the tumor single cell sequences. In particular, BEAM, SCG and SCITE performed well in imputing missing bases for datasets with a large number of cells. Our results also confirm Ross and Markowetz (2016) conclusion regarding the accuracy of SCG for datasets representing a large number of cells but containing few SNVs. However, we have newly reported that the gain in accuracy is due to correct imputation of missing data, and that SCG does not perform well in correcting FPs and FNs. In fact, all methods require a large number of SNVs to detect and correct FPs and FNs. And, as mentioned earlier, no other methods provided information on their performance in accurately imputing MBs and correcting FPs and FNs, so our results provide the knowledge of potential errors that each method may produce in actual empirical data analyses, which will be a guide to analyze the inferences of these methods for practitioners.

We have shown that, when the number of SNVs sampled is large, many methods also show good performance in detecting and correcting false positive and false negative mutation assignments. In our analyses, BEAM, SCITE, and SiFit methods performed very well for datasets containing a small number of cells, both for small and large number of SNVs. These three methods employ molecular phylogenetics, but BEAM is based exclusively on classical molecular phylogenetic methods and applies a Bayesian framework to impute MBs and correct FPs and FNs by using the initial cell sequence phylogeny as a prior. In contrast, SCITE and SiFit employ a model that account for sequencing error rates in the process of inferring the evolutionary tree of cells, which also results in improved cell sequences (Jahn, et al., 2016; Zafar, et al., 2017). SiFit produces refined cell sequences by inferring the order of mutations along the inferred maximum likelihood cell phylogeny with given error rates, whereas SCITE uses a Bayesian method to search for a cell phylogeny that intrinsically maps mutations along branches in the phylogeny. We consider BEAM and SCITE to be in the same class of methods, with the difference that SCITE needs to model sequencing error rates but BEAM does not. Interestingly, both of them show comparable results, which are among the best. That is, BEAM obviates the need to apply the same sequencing error rate model throughout the tree, which may be preferable because mutational patterns change in cells based on their evolved state (Frank and Nowak, 2004), potentially resulting in heterogeneity of error rate models among clones.

Our results show that methods that incorporate the cell phylogeny are more powerful than others, especially when the number of cells per clone is small and the number of SNVs is large. This is because BEAM, SiFit, and SCITE perform better than OncoNEM, which aims to infer clone phylogeny, and SCG, which employs a mixture model that identifies groups of cells with shared clone genotypes. Because tumor cells descend from pre-existing cells, there is evolutionary continuity in cell sequencing datasets, which enables computational methods to correctly impute missing data and make correct base assignments. We find that molecular evolutionary methods that have been successfully applied for species and strain phylogenetics for decades serve as a strong foundation for phylogenetic approaches with greater power to impute missing data and refine cell sequences for small datasets.

## Acknowledgements

We thank Heather Rowe and Chivonne Matthews for comments and editorial support. We also thank Zachary Hanson-Hart for technical support.

